# The Improvement of TSP Fertilizer Production & Quality

**DOI:** 10.1101/213223

**Authors:** N. Chaouqi, M. El Gharous, M. Bouzziri

## Abstract

Results obtained showed that some essential manufacturing factors needs to be respected throughout the production loop. These factors will help obtaining a slurry exits with the desired chemical characteristics of P_2_O_5SE_ (33%), Al (10%), P_2_O_5SE + Citrate_ (34.5%), H_2_O (22%), P_2_O_5total_ (39%)). It is necessary to keep a temperature within 100 ± 5 ° C., with a vapor pressure of 6 to 7 bars, an average density of H_3_PO_4_ (1470 g/l) and the solid rate in the feed acid less than 2%. After the reaction phase, the slurry is conveyed to the granulator where it is mixed with the recycled product at a recycling rate (RT) close to 3 and a K ratio between 0.1 and 0.2. This parameter is very important to be monitored to ensure a Good penetration of heat into the next phase in a relatively short time. During the drying process, the product humidity at the outlet must be in the range of 3.5% to 5% to allow the reaction to continue during the developing period. At the exit of the drying tube, the product is in very variable grains diameter (D), the objective of the following step is to extract the grain size range between 2 and 4 mm. To do so we changed the mesh size of the lower webs to 3/50 and 3.2 / 30 mm, which reduces the quantity of particle size less than 2%. Thus, the particles of diameter more than 3.15 mm must be continuously controlled to reach 64%. This portion will ensure conformity of the product for D_50_ and for the two intervals [2-3.15 mm] and [3.15-4 mm]. In the Storage Hall the reaction continues, the unconverted phosphate will be attacked by the unreacted H_3_PO_4_, and according to a monitoring of the evolution of the chemical composition of the final product (TSP). The required ripening time is limited to 21 days. During this study, we were able to identify and modify a number of parameters in different production phases to improve the quality of TSP according to AFCOME, knowing that each phase of the process is a client of the previous phase and supplier of the next one.

## INTRODUCTION

Phosphorus is one of the most essential elements for plant growth after nitrogen (Malakooti, 2000). Phosphorus plays an important role in many physiological and biochemical plant activities, one of the advantages of feeding plants with phosphorus is to create deeper and more abundant roots ((Arpana et al., 2002), (Mehrvarz et al., 2008)).. Phosphorus causes early maturation in plants, decreases grain mold, increases leaf chlorophyll content and may positively affect photosynthesis, thus improving crop quality (Altieri, 1995; Maene, 2000). However, the availability of this nutrient for plants is limited by different chemical reactions (formation of strong links between phosphorus with Ca^2 +^ and Mg^2+^ in alkaline pH and the same links with Fe^2 +^ and Al^3 +^ in acid soils (Chaouqi et al., 2017)) (Compaoré et al., 2001); (Malakooti, 2000). Thus, by certain agricultural practices such as the excessive application of chemical phosphate fertilizer, a large proportion of phosphorus in chemical fertilizer becomes unavailable for plants after its application in the soil (Tremblay et al., 2011). When phosphate fertilizers are not used properly, in addition to economic loss, leads to soil degradation. Therefore, better management of the fertilizer industry can be very effective in eliminating this possibility (FNCUMA, 1996). Thus, to improve the efficiency of phosphate fertilizers, they must be well distributed in the soil in order to increase the chances of contact with the roots (the 4R Stewardship, Decision-Making-Guide-Phosphorus, by IPNI-USA, 2017) (Mazoyer et al., 2017).

The production of TSP requires phosphoric acid (H_3_PO_4_), and treated rock phosphate containing respectively 42% and 30% of P_2_O_5_. In order for the TSP fertilizer to meet norms of AFCOME specifics, it needs to have a minimum chemical content of 47% P_2_O_5total_, 42% P_2_O_5SE_, 46%P_2_O_5SE+Citrate_, and a maximum of 22 ppm Cd and 2% Al. Also, characteristics in terms of particle size 1to 2mm (1- 4%), 2 to 3.15 mm (24-36%), 3.15 to 4mm (48-62%),4 to 5mm (5-8%) and D_50_ of 3.25mm ± 0.25. The objective of this study is to ensure this physicochemical quality of Moroccan TSP production. A critical analysis of the TSP production chain was initiated in the Maroc Chimie division, OCP-Safi, its objective is to increase the overall performance of the cropping systems by providing a balanced P fertilization which gives an optimal economic return. This objective, can only be achieved if the operating conditions are established strictly to AFCOME specifics, controlling the process step parameters, and making a chemical analysis of finished product.

## MATERIALS AND METHODS

### The TSP production process

The production of TSP fertilizers according to the Saint-Gobain process (Fig. 1) involves attacking the phosphate with phosphoric acid (42% P_2_O_5_).

**Figure 1:**
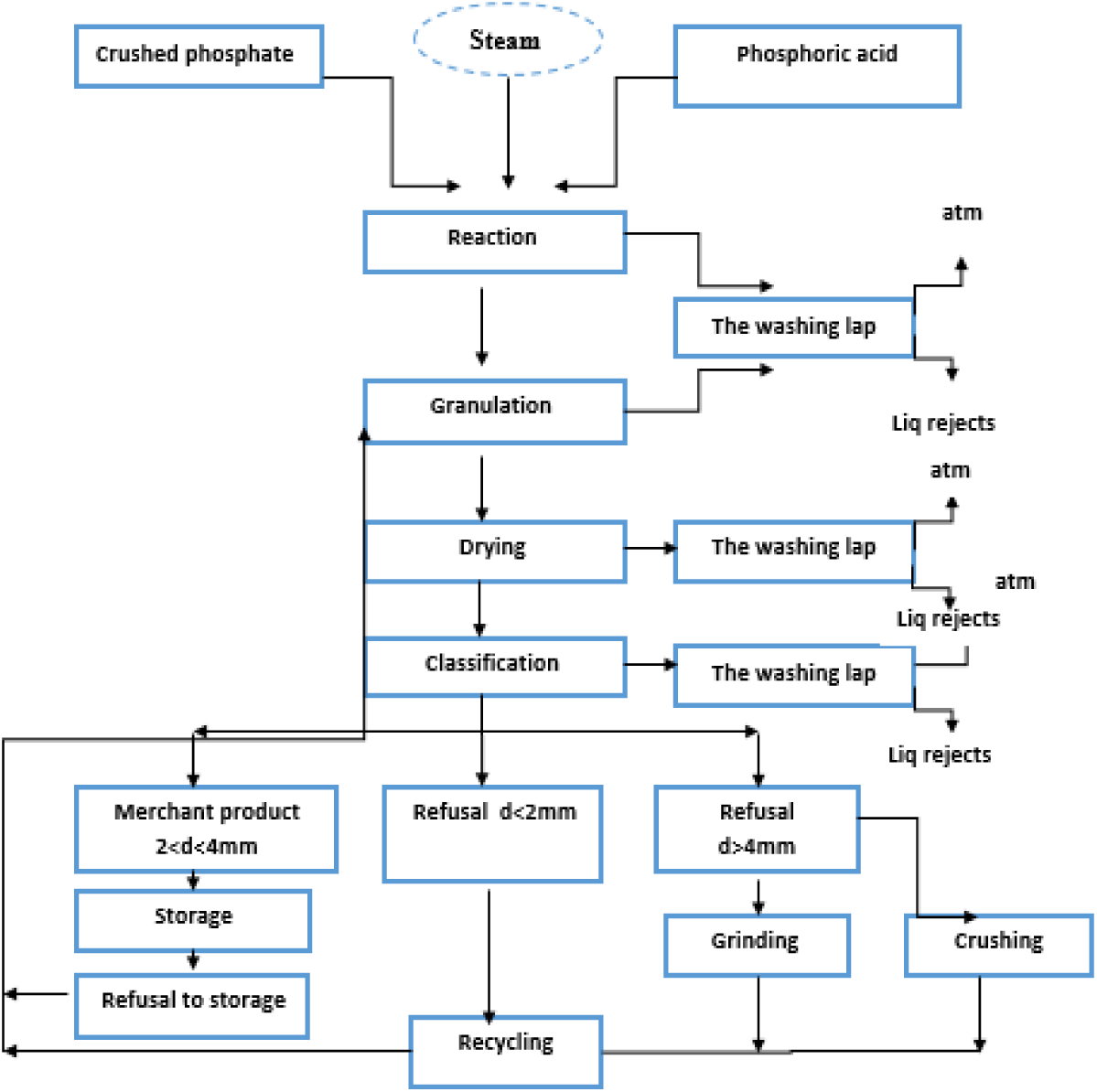
TSP production process.

The objective of this attack is to obtain mono-calcium, monohydrate phosphate soluble in water and therefore directly absorbed by plants, according to the overall reaction:

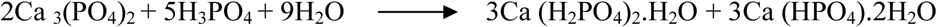

### Analytical methods used for monitoring and control of modified parameters

#### Humidity measurement

It is determined by the mass loss after drying for 4 hours at 60 ± 5°C.

#### Percentage of P_2_O_5SE&SC_ and the acid not reacted (AL)

According to the Physicochemical Laboratory Analysis of the ‘Division Maroc Chimie (OCP-Safi)’ ME00-PSC-2-ICS / P / C / L.

## RESULTS AND DISCUSSION

### Reaction

Mass flow rate of the feed tank:

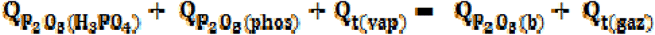

Partial flow (relative to P_2_O_5_):

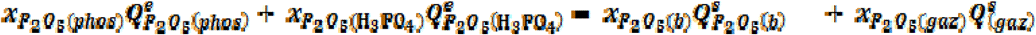

In order for slurry to obtain the chemical characteristics P_2_O_5SE_ (33%), AL (10%), P_2_O_5SE + Citrate_ (34.5%), H_2_O (22%), P_2_O_5_total (39%)), for a flow rate, Q (P_2_O_5TSP_) = 27.3 t / h and a slurry mass of 0.075t/t_TSP_, several parameters are taken into consideration:

### The particle size of phosphate feed

On a laboratory scale, analyzes show that ensuring a phosphate particle size of 90% of the past with a 160 μm sieve increases the attack surface, the chemical reaction is easier as the surface area offered to reagents is larger, requiring shorter period and less acid to be attacked.

### Acidulation rate

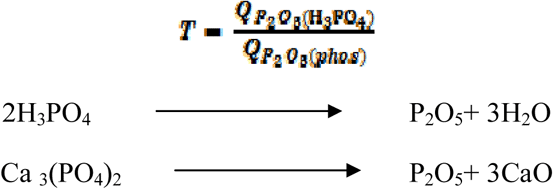

This ratio is defined as the amount in grams of P_2_O_5_ supplied by the acid required for the attack of 100 g of phosphate.

Theoretically, T = 2

Practically, this ratio varies between 2.4 and 2.6, the impurities brought by the raw materials, consume more acid.

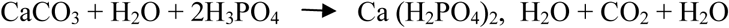

There is a consumption of H_3_PO_4_ to give slightly or not soluble phosphates. Phosphoric acid was retrograded. Also, as the carbonate content increases, the hardness and density of TSP increases, which causes granulation difficulties.

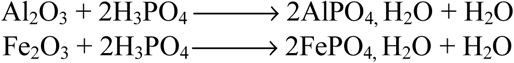

The crystalline forms of Al and Fe are insoluble in phosphoric acid. Hence an increase in the level of solid in phosphoric acid.

When the granulated phosphate rock is attacked with ortho-phosphoric acid (42%), there will theoretically be the production of phosphate mono-calcium called the triple super phosphate (TSP). In reality; this reaction is no longer complete for reasons of reagent characteristics:

The impurities present in phosphoric acid and in phosphate (Often the same quality delivered, presents an unstable profile), lead to reactions which are far from simple; they consume acid without recovering the product in the form of additional soluble P_2_O_5_.

But these two parameters depend not only on the fertilizer service, the other services require a particle size of 80 μm. As well as, for the quality of the feed material, for the H_3_PO_4_ production service: Fe_2_O_3_ has an important effect on the viscosity of the slurry. Al_2_O_3_ has a positive effect because it binds with the F^−^ and thus eliminates the negative effect of it as a highly corrosive agent. It also improves the crystallization allowing a regularity of growth of the crystals in all directions of the space. CaO has a positive impact on productivity, reacts with silica (non-reactive) and reduces its impact on the progress of production.

### Temperature (T°C)

The temperature of the tank must be between 100 and 105 °C. If T° > 105 °C this causes a bad flow of the slurry and then a bad granulation, and if T ° <100 ° C Gets a bad attack. The control is made by the steam flow through the control panel. The TSP production room uses steam about 3 to 4 t/ hour per line with a pressure of 6 to 7 bar and a temperature of 180 to 220 ° C. The rate of steam flow is regulated by a regulating valve before injection into the steam tank using steam injectors of 4 to 6 holes.

### Density (d)

To make a slurry as smooth as possible with the desired quality, a density of H_3_PO_4_ of 1480 must be ensured. A lower value (excess dilution water) causes a decrease in the temperature of the tank and the slurry becomes too fluid, the flow of phosphoric acid must be lowered. If d > 1480 (inadequate dilution water) leads to a dusty circuit, with deregulation of titers, in this case it is necessary to check the condition of the densimeter.

### Solid content (T. S)

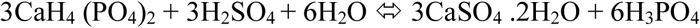

The presence of H_2_SO_4_ in the reaction mixture causes a decrease in the percentage of mono-calcium phosphate which affects the composition of the TSP, in fact a H_2_SO_4_ > 18 g/l ratio leads to a decrease of P_2_O_5SE_. Thereafter the rate of solid (T.S > 2%) produces an influence on the composition (total P_2_O_5_) of TSP. A high solids content cause a consuming difficulties of acid, acid impurities react with other species during the reaction decreasing the amount of P_2_O_5total_.

### Granulation

The mass balance:

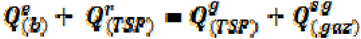

The partial balance relative to P_2_O_5_:

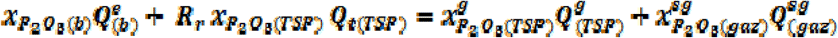

The flow rate of the granulated TSP:

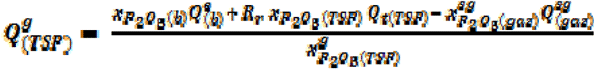

Ensuring a constant production rate is mainly based on a recycling rate constant, which depends to finished product quantity.

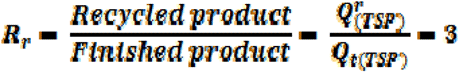

The recycling rate is between 180 and 220 T / h with a maximum capacity of 350 T / h. The product of recycling constitutes the support of granulation phase, which is in function of flow rate and slurry humidity. The granulation of the product requires in the granulator a K ratio between 0.1 and 0.2.

- If K < 0.1 the efficiency of the granulator decreases causes a dusty product.
- If K> 0.2 the granulator requires frequent stops for cleaning.

Thus, the temperature of the recycled product must be carefully controlled, to ensure a value enter 70 and 76°C.

### Drying

The control of the flow rate of the hot gases allows sufficient drying with a product leaving at a temperature of 76 °C., this T °C being reached progressively, as the product moves through the dryer. The main drying parameters having an influence on the physicochemical quality of the finished product are:

#### Product temperature

The drying temperature is limited by the melting point of the fertilizer (melting point: 190°C & decomposition temperature: 240 °C (according to the TSP FDS)), the product is admitted to the dryer at a temperature close to 85 °C. It is heated to just reach the temperature for which there is sufficient evaporation of water.

#### Dried product humidity

The TSP leaving the dryer with a humidity about 5%. The product is dried at approximately 5% to allow the further reaction between the free acid and the still unreacted phosphate to continue; this reaction cannot continue if the humidity is less than 3.5%.

#### Drying time

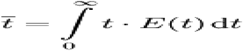

Depends mainly on the speed and inclination of the dryer tube. In fact, for a good penetration of tempture in grains in a relatively short time, the recorded rotation speed is: 3.5 r/min for a maximum capacity of: 350 T / h, with a slope of: 3 %.

### Sieving

At the exit of drying tube,the grains diameter is very variable. The objectif of this operation is to extract from the product the grain size between 2 and 4 mm. The large particles, after crushing grinding, are recycled as well as fines to granulator.

The change in sizing of 2/50 and 2.5/30 mm by 3/50 and 3.2/30 mm of the sieves reduces the fines passing quantity through the finished product,which allowed to enter in the interval [1 -2 [. The two figures below summarize the most profitable results from various tests carried out at this point. (The results displayed are those most adopted in the following months).

**Figure 2:**
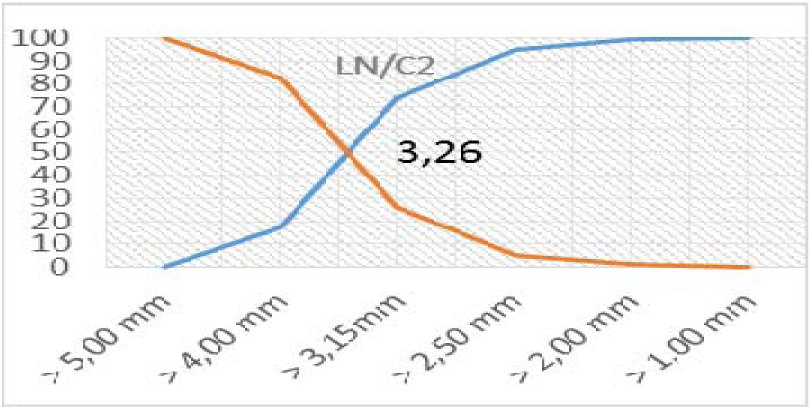
The changing size of sieves.(LN/C)

**Figure 3:**
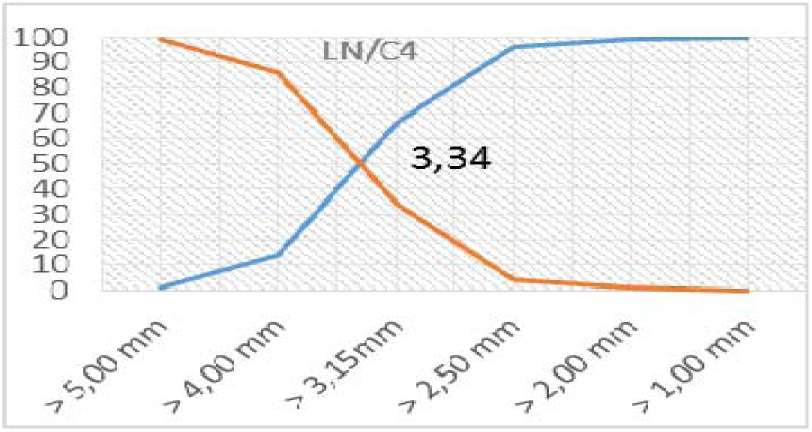
The changing size of sieves. (LN/C_4_)

In order to have a cut of 4 mm, the corresponding opening must be between 4.4 and 4.8 at the top sieve and a cut of 2 mm its corresponding opening between 2.2 and 2.4 mm for the lower sieve. It can be said that the efficiency of the two upper sieves in this case is satisfactory with more than 90%.

**Figure 4:**
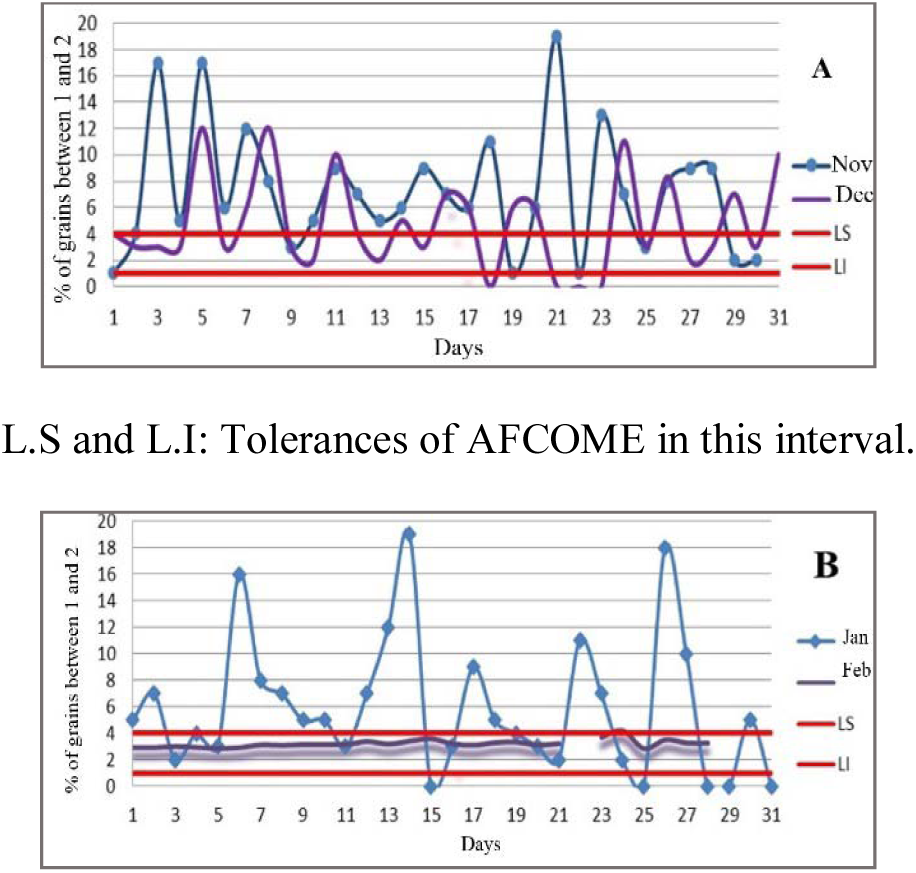
Graphical presentation of North Line before the change (A) and after (B) in [1 -2 [.

The requirements of AFCOME appear in the interval [1-2 [, this is reflected by the change of the meshes dimensioning of the lower webs. There is also a progression for the interval [4-5[due to the minimization of the mesh surfaces of the upper canvases of the sieves.

**Figure 5:**
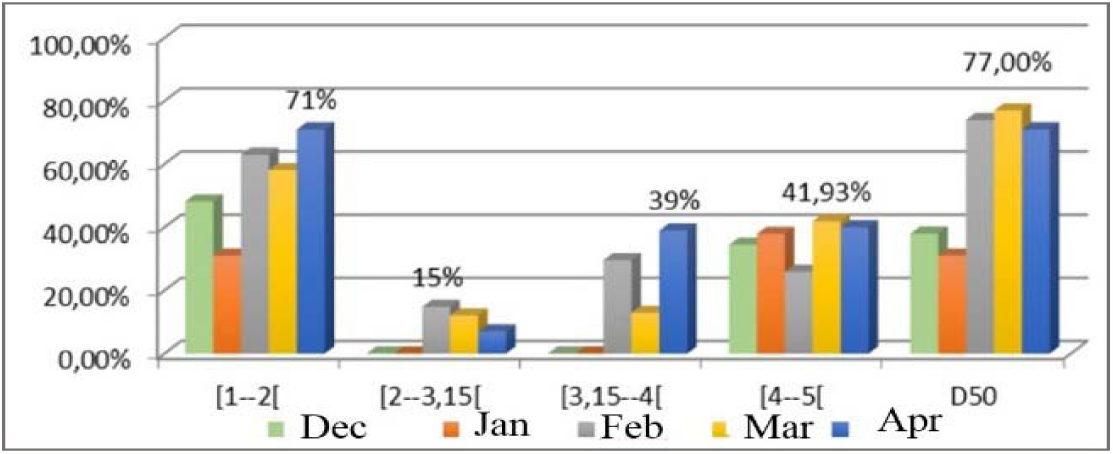
Presentation of the compliance rate according to AFCOME at the North line.

**Figure 6:**
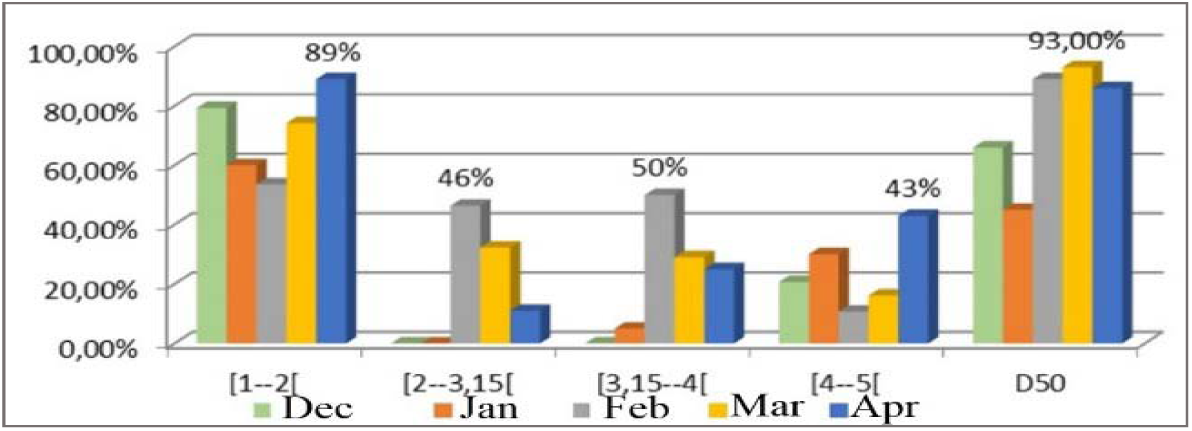
Presentation of the T.C compliance rate according to AFCOME at the South line.

These graphs result in the change of the T.C of the months before (December and January) and after (February & March & April) the change of the canvases. A remarkable evolution exceeds 70% compliance at the interval [1-2 [and median diameter 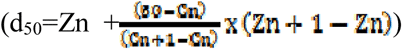 for the two production lines. Also for grains portions entering the interval [4-5[.

**Table 1:** The effect of the proportion > 3.15 on product conformity (LS). (1conform, 0 not conform)

| [3,15-4[ | [2-3,15[ | D<sub>50</sub> | > 3,15 |
| --- | --- | --- | --- |
| 1 | 1 | 1 | 72 |
| 1 | 1 | 1 | 67 |
| 1 | 1 | 1 | 64 |
| 1 | 1 | 1 | 69 |
| 1 | 1 | 1 | 63 |
| 0 | 0 | 1 | 50 |
| 0 | 0 | 1 | 59 |
| 0 | 0 | 1 | 43 |
| 0 | 0 | 0 | 34 |
| 0 | 1 | 1 | 56 |
| 0 | 0 | 1 | 51 |

**Figure 7:**
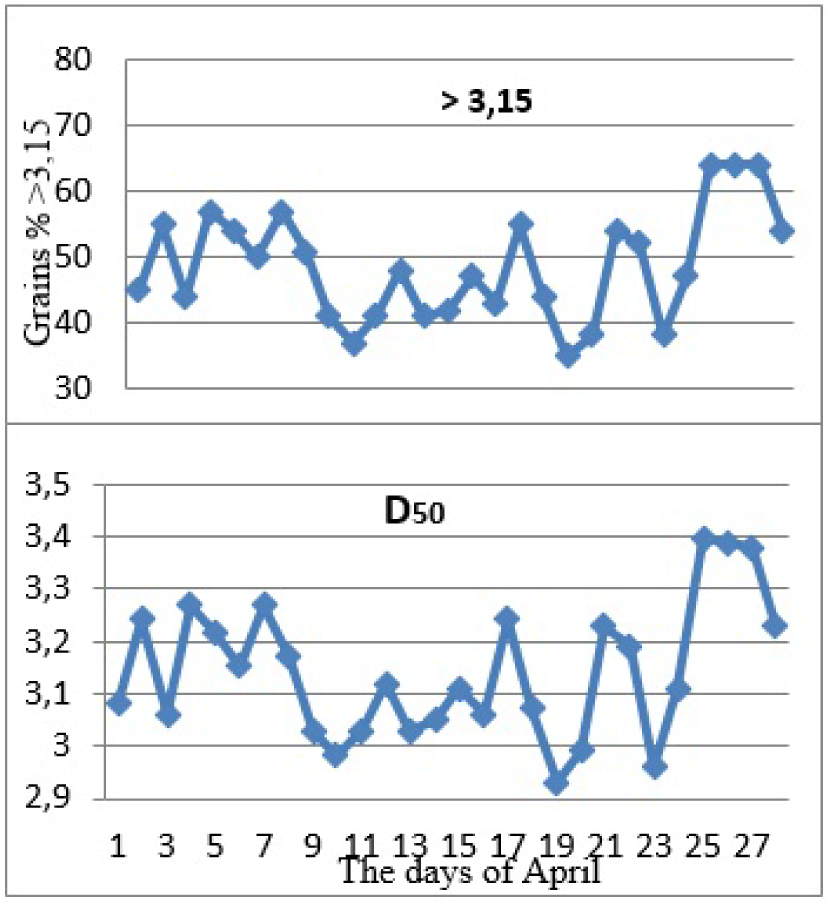
the evolution of the D_50_ f (3.15)

According to particle size analyzes results, the grains proportion with an average value of 64% of 3.15 cm diameter, is that which influences the conformity of product in the two intervals [3.15-4 [, [2-3.15]. These modifications are considered to be beneficial at the Storage Hall, namely a remarkable reduction in the refusal with 1.86%.

### Storage

In the storage hall the reaction continues, the unreacted phosphate will be attacked by the unreacted phosphoric acid, to reach the final stage of this reaction and meet both the commercial specifications (AFCOME Requirements), A storage time is required for the product, known by the ripening time. Determining of ripening time, is do it by evolution of chemical composition of TSP product (FP-TSP), namely P_2_O_5SE_, P_2_O_5SE+ SC_, total P_2_O_5_, %AL and product humidity.

From the results obtained (Tab 2) it can be said that the time required to satisfy the AFCOME requirements is 21 days. From the physical analysis (Tab 3), it can be seen that the thermal cycle in the Storage Hall (REX1) does not influence the physical quality of the TSP during this ripening period.

**Table 2:** Evolution of the chemical composition Output-Line.

| Days | %<br>P<sub>2</sub>O<sub>5SE</sub> | %<br>P<sub>2</sub>O<sub>5SE+SC</sub> | %<br>P<sub>2</sub>O<sub>5total</sub> | %<br>AL | %<br>H<sub>2</sub>O |
| --- | --- | --- | --- | --- | --- |
| 1 | 40,5 | 43,7 | 46,1 | 5,7 | 5.8 |
| 3 | 40,8 | 44,4 | 46,2 | 2,8 | 5.3 |
| 5 | 40,9 | 44,5 | 46,7 | 2,1 | 5.1 |
| 7 | 41,1 | 44,7 | 46,6 | 1,9 | 5 |
| 10 | 40,9 | 44,7 | 46,7 | 1,6 | 5.1 |
| 14 | 41,3 | 44,7 | 47,2 | 1,5 | 5 |
| 16 | 41,8 | 45,4 | 47,7 | 1,4 | 3.7 |
| 19 | 41,8 | 45,8 | 47,7 | 1,4 | 3.4 |
| **\*21** | **41,8** | **45,7** | **47,7** | **1,4** | **3.2** |
| **22** | 41,8 | 45,7 | 47,7 | 1,4 | 3.2 |
| **23...29** | 41,8 | 45,7 | 47,7 | 1,4 | 3.2 |

**Table 3:** Particle size of FP-TSP in the Hall.

| >5<br>mm | >4<br>mm | >3,15<br>mm | >2,5<br>mm | >2<br>mm | >1<br>mm | Days in<br>the hall |
| --- | --- | --- | --- | --- | --- | --- |
| 0 | 7 | 41 | 88 | 97 | 100 | Line Output |
| 0 | 7 | 40 | 88 | 97 | 100 |  |
| 0 | 6 | 38 | 86 | 96 | 100 | 7 Days |
| 0 | 5 | 35 | 84 | 95 | 100 |  |
| 0 | 6 | 43 | 88 | 97 | 100 | 12 Days |
| 0 | 8 | 37 | 86 | 97 | 100 |  |
| 0 | 8 | 44 | 90 | 98 | 100 | 18 Days |
| 1 | 6 | 41 | 87 | 96 | 100 |  |
| 0 | 8 | 37 | 86 | 97 | 100 | 21 Days |

The presentation of the compliance rate of finished product TSP during the months of February, March and April, at the level of the two storage halls:

**Figure 8:**
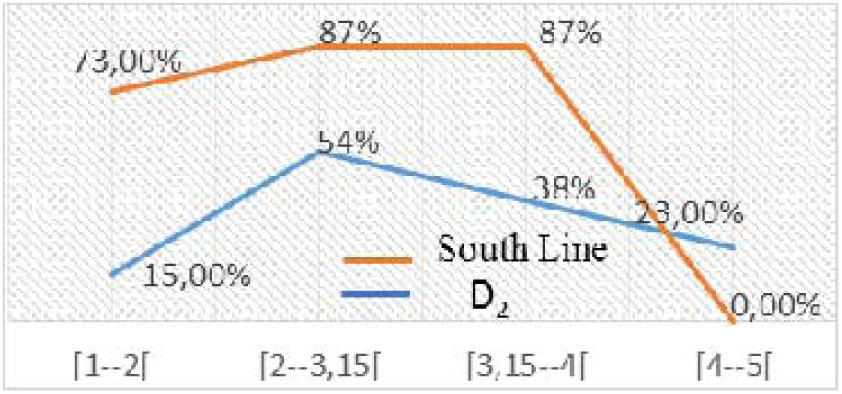
Comparison of the compliance rate between South Line & Hall-D_2_ output.

**Figure 9:**
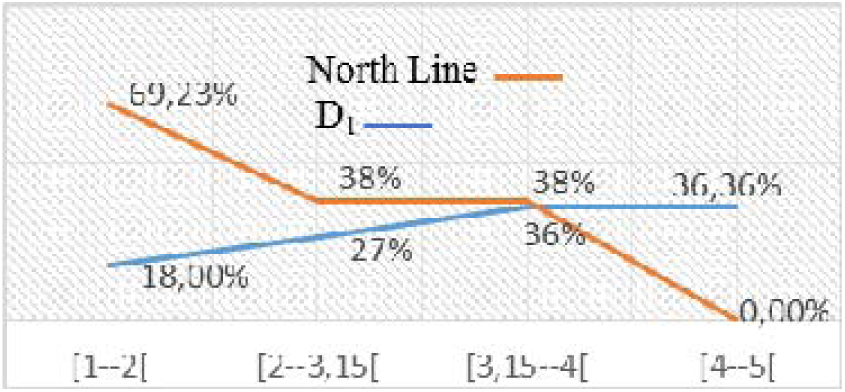
Comparison of the compliance rate between North Line & Hall-D1 output.

There is a remarkable drop in conformity of line to the destocked, this quality degradation due first of all to the mixing of conforming quality production days with others not in conformity.

This mixture affects not only the physical quality but also the chemical. To avoid this risk, we have established the FIFO strategy, the following lot will not be started until the previous lot has been exhausted. In order to respect the ripening time of the product according to the date of its arrival at the Hall.

## CONCLUSION

Fertilizers are one of the key factors for agricultural development to promote food security and maintain agricultural productivity of soils. A reasoned use of fertilizers remains the key to achieving these objectives. To produce according to AFCOME, it meets criteria such as chemical quality and physical quality, these criteria make fertilizer production more technically advantageous: A mixture that has a well-defined constitution of elements with a higher dose of principles Fertilizers, economically and socially: A considerable increase in yields per unit of cultivated area. In order to achieve the AFCOME characteristics, it is necessary to progressively apply the approach to the whole.

